# Novel methodology for localizing and studying the insect dorsal rim area morphology in 2D and 3D

**DOI:** 10.1101/2022.02.07.479135

**Authors:** Vun Wen Jie, Arttu Miettinen, Emily Baird

## Abstract

Polarized light-based navigation in insects is facilitated by a specialised polarisation-sensitive part of the eye known as the dorsal rim area (DRA). Existing methods to study the anatomy of the DRA are in most cases destructive and very time consuming. Here, we present a novel method that allows for DRA localization using 3D volumetric images acquired through micro-computed tomography in combination with 2D photographs. We used the method on size polymorphic buff-tailed bumblebees, *Bombus terrestris*. We found that the size of the DRA can be easily acquired from photographs of the dorsal part of the eye. An allometric analysis of the size of the DRA in relation to body size in *B. terrestris* showed that the DRA region increases with body size but not at the same rate. Using our method, we localised the DRA in a 3D volume rendering of the eye using 2D photos localization of an individual’s DRA onto the volumetric image stack of the head allowed for individual-level descriptions of the ommatidial structures (lens, crystalline cones, rhabdoms) to be performed on three different eye regions (DRA, non-DRA, proximate to DRA). The only feature distinctly unique to the DRA ommatidia was the smaller dimension of crystalline cones in comparison to other regions. Using our novel methodology we provide the first individual-level description of DRA ommatidial features and a comparison of how the DRA area varies with body size in bumblebees.

## 2. Introduction

Navigation is pivotal for many animals to their day-to-day tasks of foraging, finding mates, avoiding competition or predation and returning home. While on such journeys, insects maintain their bearings using directional information acquired from the position of the sun (Dyer and Dickinson, 1994; Wehner, 1989; Wiltschko and Wiltschko, 1980), moon and stars (Dacke et al., 2004, 2003a; Nørgaard et al., 2008; Papi, 2001; Wehner, 1992), the spectral light distribution across the sky (Wehner, 1997) and landmarks (Bisch-Knaden and Wehner, 2003; Bregy et al., 2008; Cartwright and Collett, 1983; Cheng et al., 2009; Knaden and Wehner, 2005; Seidl and Wehner, 2006; Steck et al., 2010, 2009; Wehner et al., 2006, 1996; Wehner, 2003, 1992). Insects can also obtain compass information using the pattern of polarization that is created in the sky (Frisch, 1967; Wehner, 1997; Wehner and Labhart, 2006) by the scattering of sunlight in the Earth’s atmosphere (Brines and Gould, 1982; Stockhammer, 1959). While this pattern is invisible to us, it can be perceived by insects through a small dorsal subsection of their compound eyes called the dorsal rim area (DRA). The DRA ommatidia – the single visual units, or facets of a compound eye – are structurally different from those in non-DRA regions and contain orthogonally arranged microvilli that enable the perception of polarized light (Goldsmith and Wehner, 1977; Hardie, 1984; Israelachvili and Wilson, 1976).

The function of the DRA has been primarily studied by analysing its structural morphology, which has now been detailed in multiple insect orders: Blattodea (Labhart and Meyer, 1999), Coleoptera (Dacke et al., 2003b, 2002, 1999; Gokan, 1990, 1989; Labhart et al., 2009, 1992; Labhart and Meyer, 1999), Diptera (Hardie et al., 1989; Labhart and Meyer, 1999; Strausfeld and Wunderer, 1985; Wada, 1974a, 1974b; Wunderer and Smola, 1982), Hymenoptera (Aepli et al., 1985; Duelli and Wehner, 1973; Greiner et al., 2007; Helversen and Edrich, 1974; Herrling, 1976; Labhart, 1986; Meyer and Domanico, 1999; Meyer and Labhart, 1981; Narendra et al., 2016, 2013; Räber, 1979; Reid, 2010; Sommer, 1979; Zeil et al., 2014), Lepidoptera (Hämmerle and Kolb, 1996; Kolb, 1985; Labhart et al., 2009; Meinecke, 1981; Stalleicken et al., 2006), Odonata (Meyer and Labhart, 1993), Orthoptera (Egelhaaf and Dambach, 1983; Homberg and Paech, 2002; Labhart and Keller, 1992). Though these studies describe the DRA morphology of a particular species, they do not provide analyses of variation between individuals (see Labhart and Keller 1992 for an exception) or complete descriptions of each of the different ommatidial components within the DRA. This is due primarily to methodological limitations – the histological and electron microscopy methods typically used in these studies are destructive and are limited to non-isotropic analyses of structures in either the longitudinal or the transverse plane in each specimen. To better understand how DRA morphology varies both within and between species, and how this might ultimately relate to differences in functionality, methodologies that are capable of describing the ommatidial features within one specimen in both 2D and 3D are necessary.

To characterize the morphological features of the DRA, one must first localize and isolate it from the non-DRA regions of the compound eye. The localization of the DRA region can be done through the identification of orthogonally arranged microvilli (Egelhaaf and Dambach, 1983; Greiner et al., 2007; Hämmerle and Kolb, 1996; Meyer and Labhart, 1993; Narendra et al., 2013; Zeil et al., 2014). In species where it has been studied, the DRA is often visually distinct from other parts of the eye. For example, in locusts with light-coloured eyes, the DRA appears as a dark region and in insects with black eyes, such as bumblebees, the DRA is often also indicated by a lighter region (Aepli et al., 1985; Greiner et al., 2007; Labhart et al., 1992; Labhart and Keller, 1992; Meyer and Labhart, 1981). Here, we describe a new method that combines 2D photography with 3D data generated from micro-computed tomography (micro-CT) to localize and characterize the DRA in eyes where it is visually distinct. Our method begins by superimposing a 2D image of the distinct DRA region of a compound eye onto the respective 3D reconstructed head model. This not only enables the localization of the DRA region that is otherwise invisible in the 3D data but also facilitates detailed descriptions of DRA morphology in individual specimens. In addition, our approach enables an analysis of how the DRA ommatidia are distributed in 3D space, which can provide insights into how they sample the skylight polarization pattern, something may have important consequences for their function within and between insect species. We applied our methodology to the bumblebee *Bombus terrestris*, which was chosen for this study because they have dark eyes with a grey dorsal region (Zeil et al., 2014) and they exhibit size-polymorphism (Goulson, 2010). We use our data to explore if and how the size of the DRA region in *B. terrestris* varies with both body and eye size. We demonstrate that we can accurately localise the DRA region by mapping the 2D images of individual eyes onto 3D volumetric scans of the same eyes and using this to guide virtual segmentation. Our approach was validated by using transmission electron microscopy (TEM) to compare the features of the virtually segmented DRA structures with features in other areas of the compound eye. Using our method, we provide the first complete description of the 3D morphological structures in DRA ommatidia in an individual insect and the first analysis of how DRA surface area changes with eye size within a species.

## 3. Material and Methods

### 3.1 Animal handling

*Bombus terrestris* colonies were acquired from a commercial supplier (Koppert, Netherlands) and kept in incubators (Panasonic MIR Cooled Incubators, 123L) at 26°C in complete darkness. They were provided with 50 % sugar water solution and fresh-frozen, organic pollen every two to three days (Naturprodukter, Raspowder Bipollen Ekologiskt bipollen).

Twenty workers of different sizes were randomly selected and assigned individual ID numbers. The selected individuals were first anesthetized with CO_2_ and then sacrificed using ethyl acetate. The thorax was photographed (Canon IXUS 220 HS, 12.1) and the inter-tegular distance (ITD), the distance between the two insertion points of the wings that can be used as a proxy for body size (Cane, 1987), was measured in pixels and converted to mm using FIJI (Version 1.8.0 _172) (Schindelin et al., 2012).

### 3.2 Acquisition of 2D DRA images and surface area measurements

A marked individual was placed in a 3D printed sample holder attached to the end of a micromanipulator (Figure 1 A). The dorsal region of the left compound eye was viewed under a stereomicroscope with a camera attachment (Figure 1 B). After removing the pile (soft hair) around the ocelli using micro-scissors, a small grid paper for size calibration was attached onto the midsection of the head. The grid was attached using super glue leaving at least one ocellus uncovered. The DRA was illuminated using a light source that was positioned at a 45° with respect to the DRA position (Figure 1 C & D).

**Figure 1:**
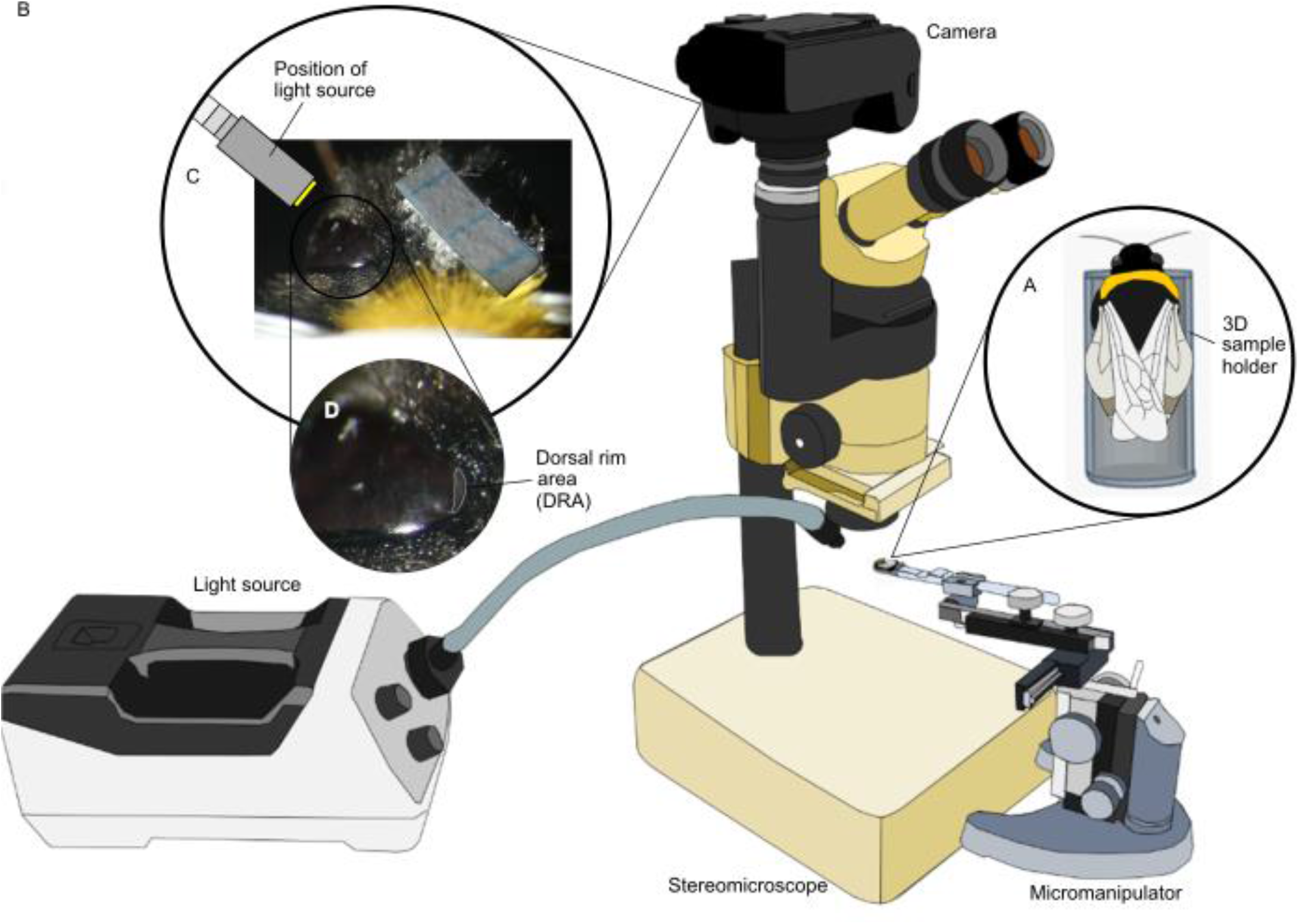
Setup and equipment for the dorsal rim area (DRA) image acquisition in 2D. (A) Bumblebee in sample holder, (B) Microscope set-up to acquire DRA photographs, (C) light set-up for DRA visualization, (D) enlarged view of the DRA.

The DRA and the grid paper were photographed (Canon EOS 70D) with a resolution of 1824 × 2432 pixels. The DRA surface area in 2D was calculated from the images using FIJI (Version 1.8.0_172) (Schindelin et al., 2012).

### 3.3 Staining and embedding

The heads were dissected from the body and a microscalpel was used to remove the mouthparts. The heads were then immediately placed in 70 % ethanol and 0.5 % phosphotungstic acid (PTA) solution for 10 days (Smith et al., 2016). The heads underwent dehydration using a graded ethanol series and were cleared in acetone prior to transferring them to epoxy resin (Agar 100). Samples in the wet resin were transferred onto a perspex block and cured in an oven at 60°C for 48 h. After curing, excess external resin was removed to expose the eyes and surrounding cuticle (Taylor et al., 2016).

### 3.4 Micro-CT

Micro-CT scanning was performed at the TOMCAT beamline of the Swiss Light Source, Paul Scherrer Institute, Villigen (Switzerland) (beamtime number 20190641). Only 14 out of the selected 20 heads were scanned since the remaining 6 were damaged during staining and embedding. The intact heads were scanned using a monochromatic X-ray beam (20 keV), 2001 projection images of 2560 × 2160 pixels with a 60 ms exposure time were collected by a PCO.Edge 5.5 detector over 180 deg and a propagation distance of 100 mm. The scans were done using a 4x objective. The projection images were processed with the Paganin phase-retrieval method (Paganin et al., 2002) and reconstructed into 3D volumes using the gridrec algorithm (Marone and Stampanoni, 2012) with Parzen filter, resulting in 16-bit volume images that had an isotropic voxel size of 1.625 µm.

The regions of interest (ROIs) from the reconstructions were cropped using the program Drishti Paint (Limaye, 2012) and resaved as 16-bit TIFF files to reduce file size. Two ROIs were obtained from the reconstructions of each sample, and they consisted of (i) the left compound eyes (in the animal’s perspective), and (ii) the dorsal head (the dorsal part of the left compound eye and the ocelli, which were used as landmarks to facilitate mapping the 2D images in later analysis, section 3.6).

### 3.5 3D Compound eye and surface area measurements

3D models were generated from the cropped compound eyes in Amira (Version 2020.3.1, ThermoFisher Scientific) using a combination of segmenting tools that included thresholding, brush, fill, shrink and grow. The surface area of the compound eye was obtained by generating a surface from their segmented label (Taylor et al., 2019).

### 3.6 Localizing the DRA on a 3D head model

From the 14 dorsal head volumes, six with best-preserved eyes and high contrast micro-CT scans were chosen for DRA localization and anatomical ommatidial analysis. We chose two samples from each of the three chosen body size ranges based on the ITD, small (2.965 mm - 3.902 mm), medium (3.902 mm - 4.839 mm) and large (4.839 mm - 5.776 mm). 3D models of the eyes were generated in Amira using a combination of segmenting tools that included thresholding, brush, fill, shrink and grow. Because the grey DRA region is not visible in the 3D volumes, the localization of the DRA of an individual bumblebee requires both its 3D dorsal head and its 2D DRA image.

The DRA was localized by overlaying a 2D DRA image (Figure 1 C) onto the 3D dorsal head volume taken from the same specimen (Figure 2 A). The ocelli were used as landmarks for precise mapping (Figure 2 B). Mapping was performed in the project view of Amira using modules that included surface view and clipping plane. The DRA on the 3D model was segmented using the mapped image as a reference (Figure 2 C). The surface area of the DRA was obtained by generating a surface using the segmented DRA labels.

**Figure 2:**
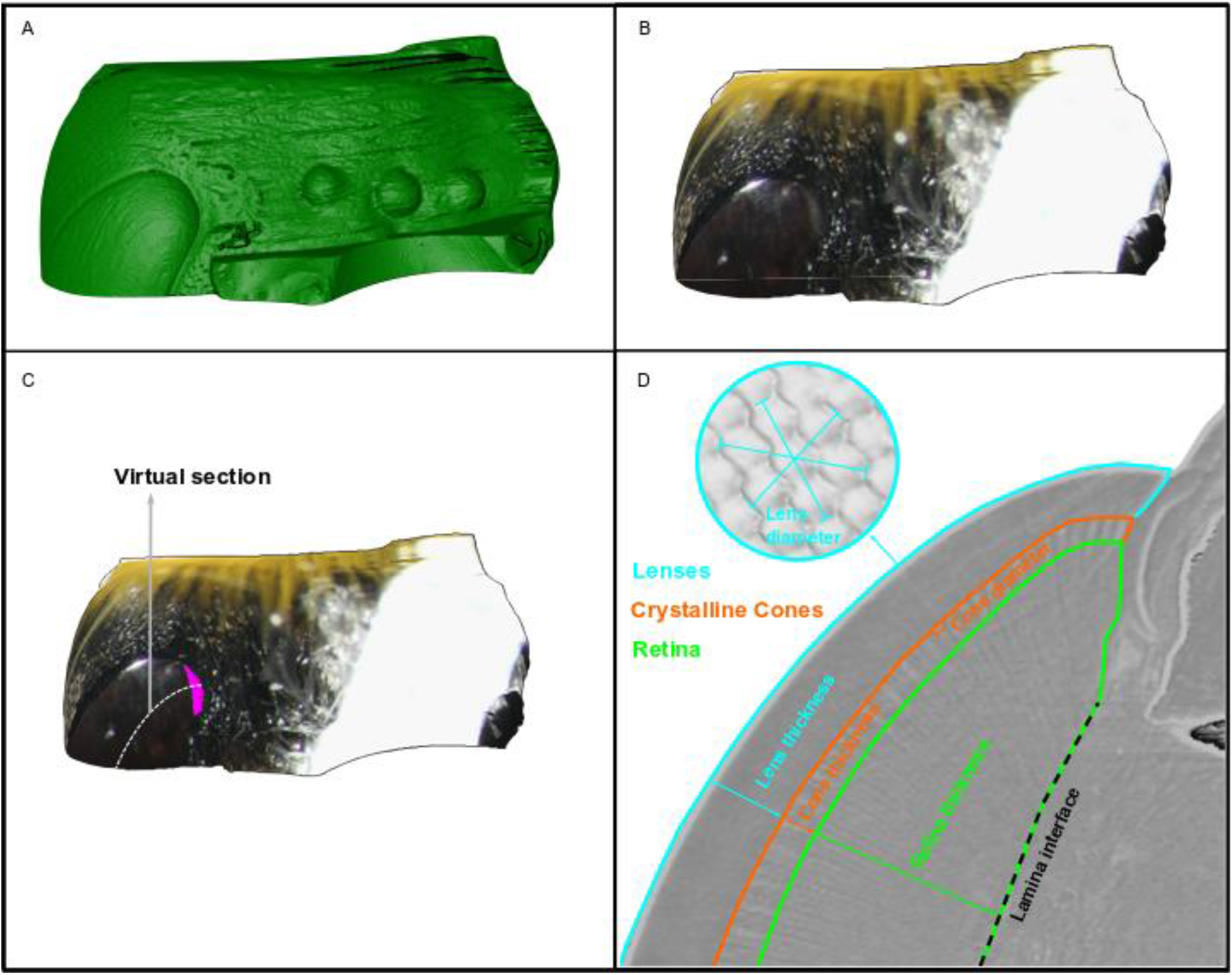
The process for localizing the DRA on a 3D head model of a bumblebee *Bombus terrestris*. (A) A 3D volume rendering of a bumblebee head. (B) A 2D photo is overlayed onto the 3D volume, (C) The overlaid image is used to identify and segment the DRA (purple outline) in Amira. A grey dotted line indicates the region where virtual sectioning is performed. (D) Virtual slice showing ommatidial ommatidial structure measurements.

To determine the accuracy of our DRA localization method, we took transverse sections of the rhabdoms in regions predicted to be DRA and non-DRA on a separate individual prepared and visualised them using transmission electron microscopy (Romell et al., 2021).

### 3.7 DRA allometry and overall compound eye surface area

Since the two methods used for determining the DRA surface area yielded very similar results (Figure 3), the surface area measurements of the DRA from the 2D photos were used for the allometric analysis of how DRA surface area varies with compound eye surface area (n = 20). The surface area of the compound eyes was obtained from the 3D compound eye models (n = 14). All variables were converted to linear measurements before logarithmic transformation. The relationships between the surface area of the DRA and compound eyes were plotted against ITD onto a log-log plot.

**Figure 3:**
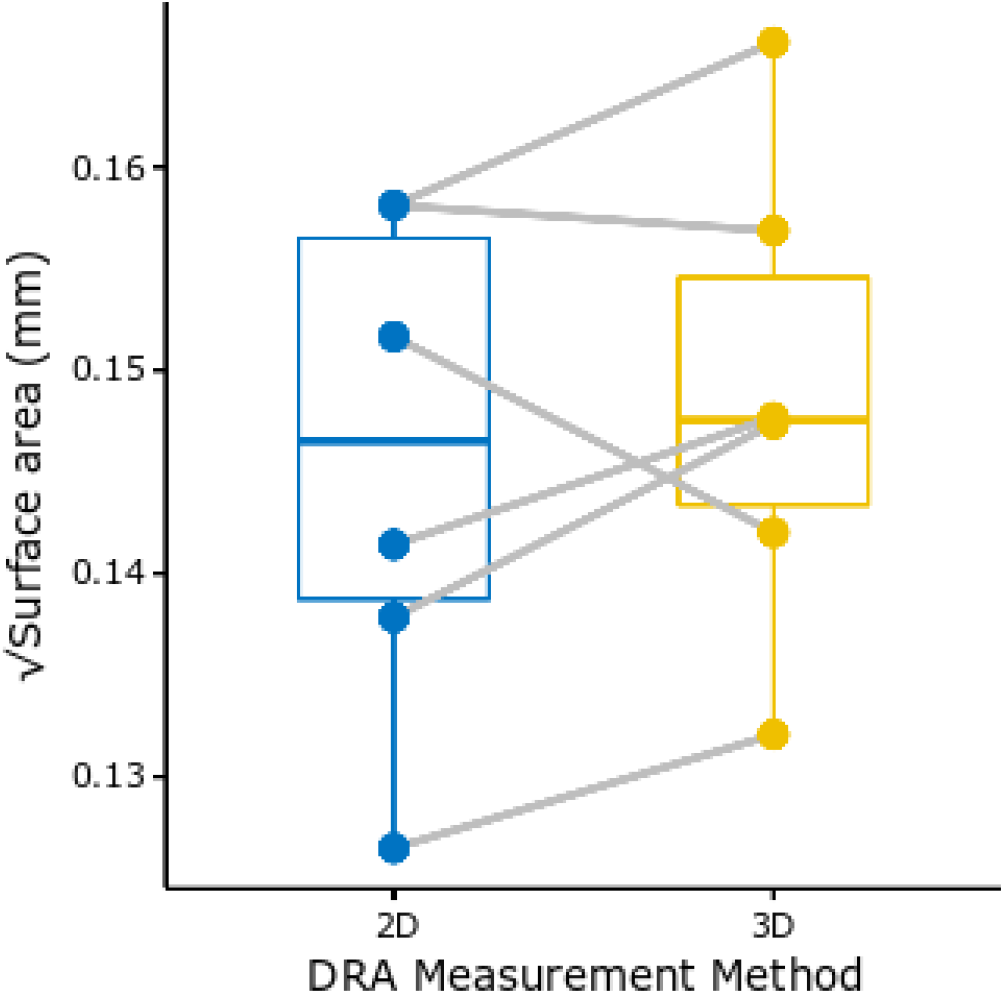
Boxplot comparing 2D and 3D DRA surface area measurements taken from the same individuals (grey lines) of *Bombus terrestris*. There was no significant difference between the values acquired using the 2D images or 3D volumes (t = −1.0421, p = 0.3451, n = 6, Wilcoxon signed-rank test).

### 3.8 DRA ommatidial structure measurements

The compound eyes from the six individuals that had their DRA localized and segmented were also used for volumetric analysis. The three chosen regions for ommatidial structure comparisons were within the DRA, proximate to the DRA (five ommatidial rows surrounding the DRA), and non-DRA (> 20th ommatidial row vertically from the DRA). Seven longitudinal virtual histological slices consisting of the three regions were obtained per eye from the 3D scans using the clipping plane module in Amira. Only virtual slices where the rhabdoms were continuous were used for further analysis (Figure 2 D). Virtual histological slices were exported and analyzed in FIJI (Version 1.8.0 _172) (Schindelin et al., 2012). The thickness of the lens, crystalline cones and rhabdoms as well as the width of the crystalline cones were measured. The lens width was measured externally on the model using 3D line tools (Figure 2 D). For each of the six eye samples, 20 ommatidial structure measurements were performed.

## 4. Results and Discussion

Our methodology allowed for the localization of the DRA on a homogeneous eye surface through the mapping of 2D images onto 3D volumetric data in individual compound eye samples and a comprehensive study of the gross internal and external morphology of the DRA structures. We discovered that external 2D images are sufficient for calculating the DRA surface area, confirming that this is a valid method for more accessible high-throughput analyses of DRAs within and between species.

### 4.1 DRA localization on a homogeneous eye surface

This is the first localization of the DRA through the mapping of 2D images onto 3D volumetric data generated from micro-CT scans. We used TEM to analyze the microvilli of transverse sections of DRA and non-DRA regions of the compound eye. The presence of orthogonally arranged microvilli in the region identified as the DRA suggests that our localization was accurate (Figure 4 A - B). In addition, since the structures are relatively flat, the comparative analysis between these and the 2D images showed that there are no significant differences between the method (t = −1.0421, p = 0.3451, n = 6, Wilcoxon signed-rank test) (Figure 3). This means that the 2D external images alone can provide an easy and high-throughput estimation of DRA surface area. Although the combination of photos and micro-CT data is not new (Ijiri et al., 2018), we demonstrate here that it can also be used to make quantitative measurements. This methodology opens up new ways to study the DRA and other external features since it allows for individual-level gross morphological descriptions of these structures.

**Figure 4:**
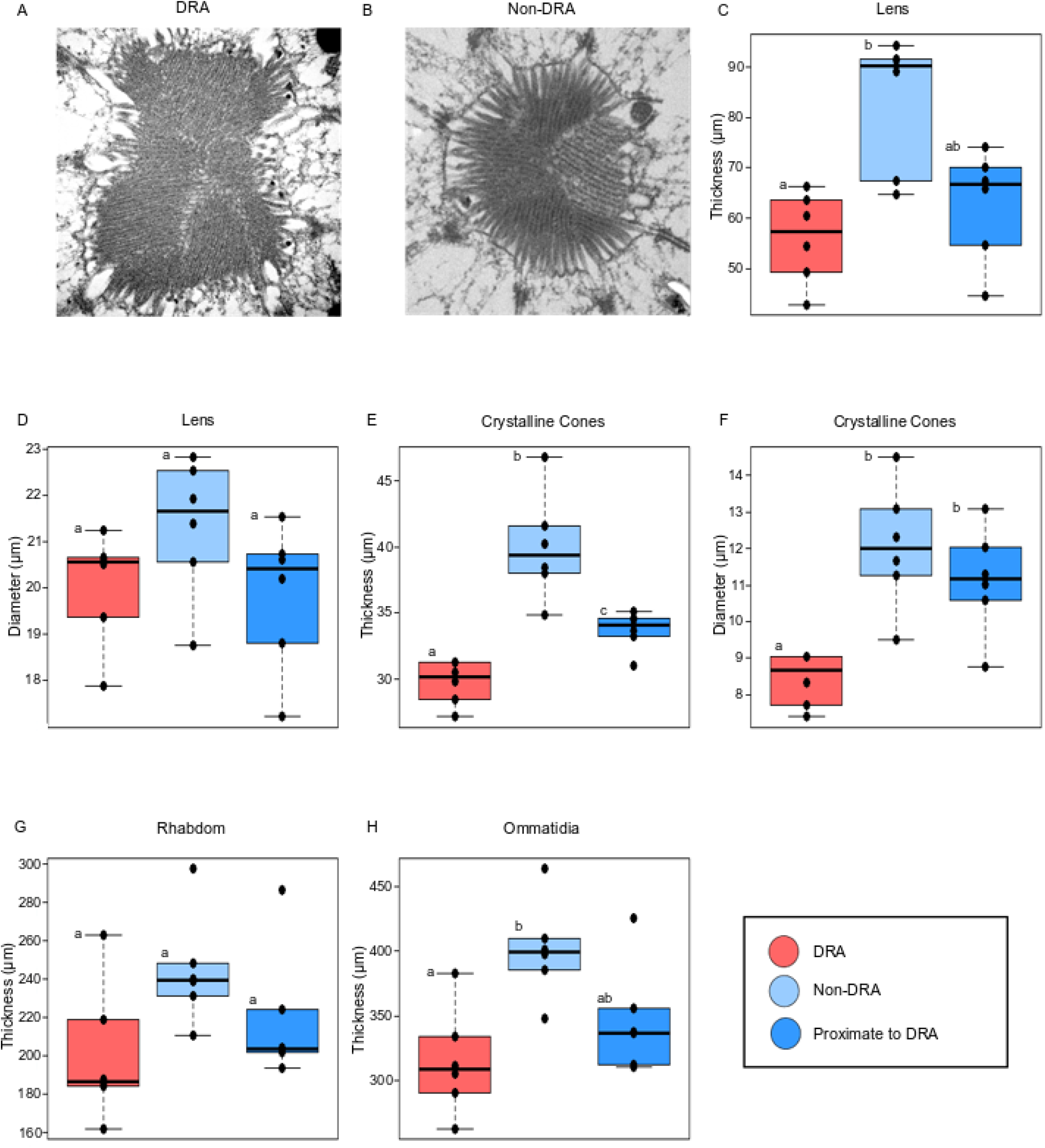
Transverse section of (A) DRA, and (B non-DRA microvilli (n = 1). Ommatidial structure comparison between DRA, proximate to the DRA and non-DRA regions. Thickness of lens (C), crystalline cones (E), rhabdom (G), and ommatidia (H). Diameter of lens (D), and crystalline cones (F) (n = 6). Letters represent the outcome of pairwise Wilcoxon Rank Sum tests, matching letters indicate no significant difference at p = 0.05 and non-matching letters indicate significance below this value.

### 4.2 A comprehensive study of the internal and external morphology of DRA structures

In contrast to previous methodology for characterising ommatidial structures in the DRA using TEM, our method allowed us to achieve gross morphological descriptions of most components of the DRA ommatidia within an individual insect due to its non-destructive nature. We found that the ommatidial structures are similar between areas within and proximate to DRA regions (Figure 4). The most distinct difference that we observed between the DRA and other eye regions was the thickness of crystalline cones; those in the DRA are both thinner (DRA vs proximate to DRA: H_2_ = 10.211, p = 0.0223; DRA vs non-DRA: H_2_ = 10.211, p = 0.0065, n = 6, Pairwise Wilcoxon Rank Sum) and smaller in diameter (DRA vs proximate to DRA, H_2_ = 13.93, p = 0.0087; DRA vs non-DRA: H_2_ = 13.93, p = 0.0065, n = 6, Pairwise Wilcoxon Rank Sum) than those in other regions. This is unlike what has been observed in other insects, such as the Canarian tiny cricket (Egelhaaf and Dambach, 1983) and the desert locust (Homberg and Paech, 2002). The question arises to as why some insects have distinct differences between the DRA and regions proximate to it while others do not? The comprehensiveness of this methodology allowed us to pick up minute differences between regions of interest and showed that structures within the DRA and proximate to the DRA can be very similar, at least in *Bombus terrestris*.

### 4.3 Method implementation on *Bombus terrestris* to study how the DRA varies with body size

This study provided an easy and high-throughput methodology for studying the DRA surface area in insects that allowed us to perform the first allometric study of how surface area of the DRA scales with body size in bumblebees. Our results revealed that the DRA, as a small component of the compound eye, correlates positively with body size (t_12_ = 2.682, p = 0.0152, n = 20, regression analysis) (Figure 5). The compound eye also correlates positively with body size (t_12_ = 7.137, p < 0.001, n = 20, regression analysis). This result is consistent with the findings of Taylor et al. (Taylor et al., 2019) and suggests that the variation of eye size with body size likely affects visual capabilities such as resolution and sensitivity in *B. terrestris*. This, in turn, is likely to affect visually guided behavior, such as the timing of activity (Spaethe and Chittka, 2003; Spaethe and Weidenmüller, 2002). Do differences in DRA size between individuals observed here also relate to functional differences, for example in navigational capabilities (by affecting the sensitivity to polarised light in the DRA)? Our method allows questions like this to be answered by facilitating the quantification of the effect of body size on the size of the DRA and could ultimately inspire more questions about DRA functionality that can be tested behaviourally.

**Figure 5:**
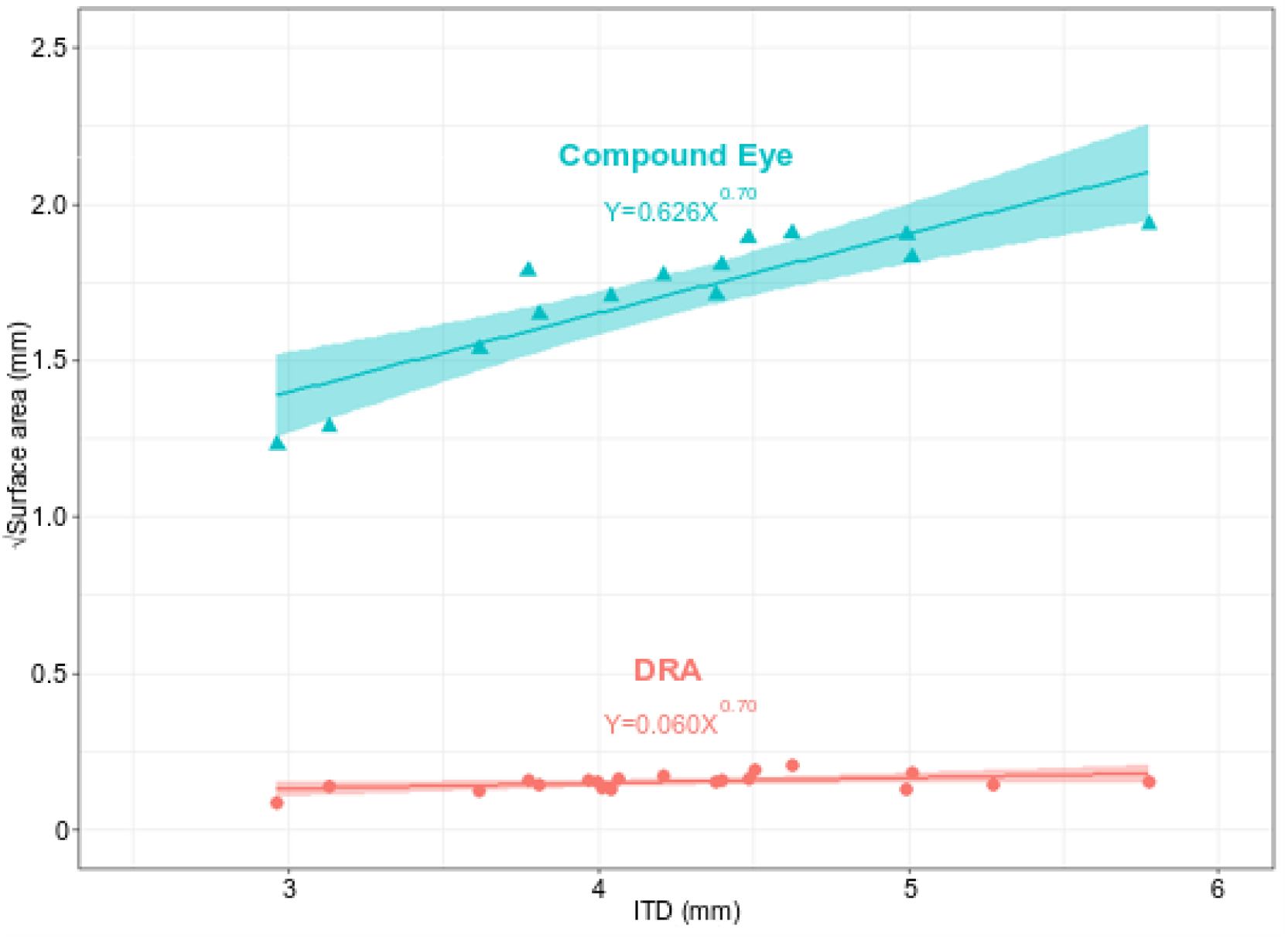
Allometric relationship between the square root of eye surface area (mm) with ITD (mm). Triangle = 3D compound eye measurements (t_12_ = 2.682, p = 0.0152, n = 14, regression analysis), Circle = 2D DRA measurements (t_12_ = 7.137, p < 0.001, n = 20, regression analysis).

### 4.4 Limitations of the method

While our method provides new insights into the morphological structures of the DRA, both externally and internally within one individual, it has some limitations that are important to highlight. Although micro-CT based 3D volume renderings of compound eyes allow for a more comprehensive gross morphological description of the DRA structure, it does not have the resolution necessary for resolving the orientation of individual microvilli. This limitation should be addressed in the future as microvilli orientations within the DRA determine how the polarisation pattern of the sky is sampled. However, as the samples prepared for micro-CT are embedded in resin and stained with heavy metal they are also appropriate for performing TEM (Handschuh et al., 2013). This correlative approach would allow for the mapping of the microvilli arrangement at the individual-level that, to our knowledge, has never been achieved in 3D. A further limitation of our method is that the 3D methodology takes a long time to learn, requires advanced amounts of computing power (gigabytes of RAM), storage space (data range in gigabytes per dataset), and can be expensive. The 2D methodology, though it is considered high throughput only works if the DRA structures are relatively flat, which is the case in *Bombus terrestris*. This is not the case in butterflies as their DRAs are curved (ranging from the edge of the eye next to the antennae to the dorsal edge of the eye) (Stalleicken et al., 2006). To acquire the surface measurements of these, the 3D localization method is necessary. Finally, this method is limited to analysing the DRAs of insects with dark eyes and whose DRA is marked by a grey region.

## 5 Conclusion

To understand insect navigational behaviour, it is necessary to study all sensory structures involved, including the DRA that is linked to polarization vision-based navigation. Though this structure is widespread among insects, it has proven to be problematic to study, due to lack of complete and non-destructive individual-level methods. Our novel methodology presented in this study allows us to produce more insightful observations of DRA morphology and, in turn, raise questions about polarization vision-based navigation in insects.

## 6 Acknowledgement

We would like to thank Inga Tuminaite for assistance throughout the project and for critically reading the manuscript drafts. We would also like to acknowledge the Paul Scherrer Institut, Villigen, Switzerland for provision of synchrotron radiation beamtime at the TOMCAT beamline X02DA of the SLS. Pierre Tichit for his technical guidance in Amira. Julia Meneghello for helping in segmenting the heads and eye surface area of the bumblebees.

